# Genomic insights into clonal diversity in UK populations of the Potato aphid, *Macrosiphum euphorbiae*

**DOI:** 10.1101/2023.05.28.542558

**Authors:** Mark Whitehead, Alison Karley, Alistair Darby

## Abstract

The potato aphid *Macrosiphum euphorbiae* is one of many polyphagous crop pests involved in the transmission of insect-vectored pathogens. While their North American counterparts reproduce via cyclical parthenogenesis, UK populations of *M. euphorbiae appear* to persist asexually, resulting in the maintenance of several genotypes, with some demonstrating genotype-specific traits; this includes innate resistance to parasitism from the hymenopterous parasitoid wasp *Aphidius ervi*. The genetic and molecular basis for genotype-specific traits is often unknown. Here we present a chromosome scale assembly for a parasitoid-resistant clonal line of *M. euphorbiae* and provide insights into the genotypic composition and distribution of UK potato aphid populations using microsatellite and whole-genome sequencing (WGS) techniques, focusing on geographically separated potato crops within two distinct areas of the UK (Merseyside and Tayside). We show that the genome consists of five chromosomal blocks, has a total size of 560 Mbp and is highly complete based on BUSCO (C: 95.6%). The sampled potato aphid populations were dominated by two genotypes, one of which is absent from commercial farm settings, suggesting either an intolerance to farming practices, such as insecticide use, or a broader host range. We suggest some putative gene functions using WGS data to explain the observed frequency of aphid genotypes. WGS data highlighted the asexual clonal lifestyle of *M. euphorbiae genotypes* in the UK, resolving individuals to a higher resolution than using microsatellite data. The work presented here will provide useful information for integrated pest management of potato aphids, elaborating on the relationship between genotype diversity and functional traits such as parasitism and insecticide resistance, and host plant use, as well as providing more resources for further comparative genomics studies within the Aphididae.

## Introduction

Aphids are important agricultural pest insects worldwide as they transmit insect-vectored viruses, causing plant disease that can result in plant disfiguration, reduced growth and ultimately a reduction in crop yield. Aphid populations can show great variation in genotypic composition and genetically-controlled phenotypic traits. An important example is the occurrence of insecticide-resistant genotypes which can dominate aphid populations when insecticide use creates a strong selection pressure (Singh *et al*., 2021). Studying the genetic structure of aphid populations can, therefore, reveal factors that underpin genotypic differences in aphid fitness, which in turn explain the ability of aphid genotypes to persist and disperse. Aphid genotypes can often show distinct spatial distributions due to physical barriers preventing spread (Guillemaud *et al*., 2003; Raboudi *et al*., 2005; Wang et al., 2016). In the case of *Aphis gossypii* (Glover), spatial differences in population genetic structure are observed even over short distances (< 1 Km) between crops in individual greenhouses (Fuller *et al*., 1999). In addition, host plant range can also play a large role in aphid spread and frequency. For example, certain *Acyrthosiphon pisum* (Harris) (Pea aphid) genotypes show greater fitness on alfalfa (*Medicago sativa*) or red clover (*Trifolium pretense*) in the wild (Via, 1991; Ferrari et al., 2006), and geographic differences in host plant availability could influence genotype spread. In sexually reproducing aphid species, host plant specialisation may drive long term speciation by reducing geneflow between these populations (Ferrari et al., 2006).

The potato aphid, *Macrosiphum euphorbiae* (Thomas), typically causes crop damage as a vector of multiple plant viruses affecting potato, including potato virus Y and potato leaf roll virus, as well as viruses that affect crops such as beans, sugar beet, sugarcane and lettuce (Blackman & Eastop, 2000). The transmission of Potyviridae viruses is especially damaging to potato crops as increased virus prevalence leads to rejection of crops grown for seed and can result in lower potato crop yield (SASA, 2017). The peach-potato aphid *Myzus persicae* is considered the most efficient virus vector on potato crops and, therefore, the major contributor to yield loss and crop damage compared with other aphid species on potato. *Macrosiphum euphorbiae*, however, still has the potential to reduce yield on certain potato varieties through virus transmission and through reduced carbohydrate accumulation and leaf rolling (Veen, 1985) when infestation levels are high (although this is infrequent in the UK: Parker, 2005).

The potato aphid is a highly polyphagous insect pest, capable of surviving on over 200 plant species, covering more than 20 taxonomic families (Blackman & Eastop, 2000; Srinivasan & Alvarez, 2010). In North America, *M. euphorbiae* is holocyclic (Simon *et al*., 2002), undergoing a sexual phase in the autumn and overwintering as eggs (MacGillivray & Anderson, 1964), using members of the Rosaceae family as primary plant hosts. Alternating between sexual and asexual forms is a common life history strategy for many aphid species, but some species persist asexually through the winter (anholocyclic lifestyle) (Dixon, 1973; Simon *et al*., 1999). Previous evidence suggests that European populations of *M. euphorbiae* persist via clonal reproduction over winter, with much rarer occurrences of sexual reproduction (Raboudi *et al*., 2012). This reproductive dimorphism is observed in other aphid species, with the asexual mode of reproduction linked to milder climates (Vorburger et al., 2003, Sandrock *et al*., 2011).

Potato aphid populations in the UK have been shown to comprise several genotypes, some of which show genotype-specific fitness traits (Clarke *et al*., 2016; Karley et al., 2017; Clarke et al., 2018). Those of specific interest include innate resistance to parasitism by the natural enemy *Aphidius ervi* (Clarke *et al*., 2017), where parasitism has a higher chance of failure in one aphid genotype (named genotype 1). Studies of the pea aphid have detected aphid genotypic differences in *A. ervi resistance* (Li *et al*., 2002; Martinez *et al*., 2014), but the genetic basis is not known. If warming climate creates more favourable conditions for asexually reproducing aphids like *M. euphorbiae*, and as pesticides become less effective or even withdrawn (Plant Health Centre, 2021), it is increasingly important to understand the prevalence and persistence of aphid genotypes that might be resistant to biological (and chemical) control (Foster et al., 2002; Clarke et al., 2018).

Here, we aim to provide a broad understanding of the genetic structure of UK potato aphid populations and more detailed information about the genetic basis for parasitism resistance. The study objectives were: i) generate whole genome sequence data for *M. euphorbiae;* ii) sample *M. euphorbiae* from potato crops over a three-year period to assess genotype frequency using microsatellite markers and whole genome sequencing; and iii) compare genome sequence data from different aphid genotypes to identify candidate genes and gene functions underpinning genotype frequency and phenotype. We report the first haploid chromosome-level genome for *M. euphorbiae* using long-range sequencing techniques, followed by chromosome-scale assembly and orientation using Hi-C and highlight possible gene functions attributed to parasitism resistance and host plant range. The data generated by our study will permit future investigations into genotype specific traits of *M. euphorbiae*, and provides a genomic resource to support integrated pest management of this crop pest.

## Results and Discussion

### Generation of genomics resources for *M. euphorbiae*

To enable the accurate assessment of population genetic structure in this study, we generated a chromosome-level genomic assembly of 50 individuals of *M. euphorbiae* of the clonal line MW16/67, which shows resistance to parasitism by the hymenopterous parasitoid wasp *Aphidius ervi* (Clarke et al., 2016). The initial Canu contig assembly, with 63 Gbp of PacBio contiguous long-read (CLR) sequel data, was 970 Mbp, close to the predicted diploid genome size (supplementary table 1). Two rounds of Haplomerger reduced the genome size to 560.8 Mbp and the total duplicated BUSCO genes to <3%. GenomeScope Kmer profiling with the Illumina data estimated a haploid genome size of 527 Mbp, in line with flow cytometry prediction of the *M. euphorbiae* genome size (Wenger *et al*., 2017) of 530 Mbp.

**Table 1.** Summary of significant GO terms with a role in host plant range and preference between *M. euphorbiae* clonal lineages. Class refers to GO terms for “Biological processes”and “Molecular function”.

| *Class* | *GO term ID* | *Name* | *TopGO P value* |
| --- | --- | --- | --- |
| BP | GO:0007608 | “sensory perception of smell” | 0.00033 |
| BP | GO:0055114 | “(obsolete) oxidation-reduction process” | 2.80E-06 |
| MF | GO:0004984 | “olfactory receptor activity” | 5.70E-05 |
| MF | GO:0005549 | “odorant binding” | 0.00035 |
| MF | GO:0016705 | “oxidoreductase activity” | 0.00154 |

After scaffolding and chromosome assignment with 10x linked reads and Dovetail Hi-C, the final scaffolded assembly showed five chromosomal linkage groups, in agreement with a previous karyotype assessment of *M. euphorbiae* (Monti *et al*., 2011) (figure 1A and 1C). The assembly consisted of 1,914 scaffolds, with an N50 of 107.7 Mbp and a total size of 560.3 Mbp, with 95.6% complete BUSCO genes (1306/1,367 insecta_odb10 database) with 86 Mbp of the contigs belonging to alternate haplotypes or repeats too divergent to collapse with HaploMerger. Thus, the MW16/67 assembly represents a near complete and highly contiguous assembly.

**Figure 1.**
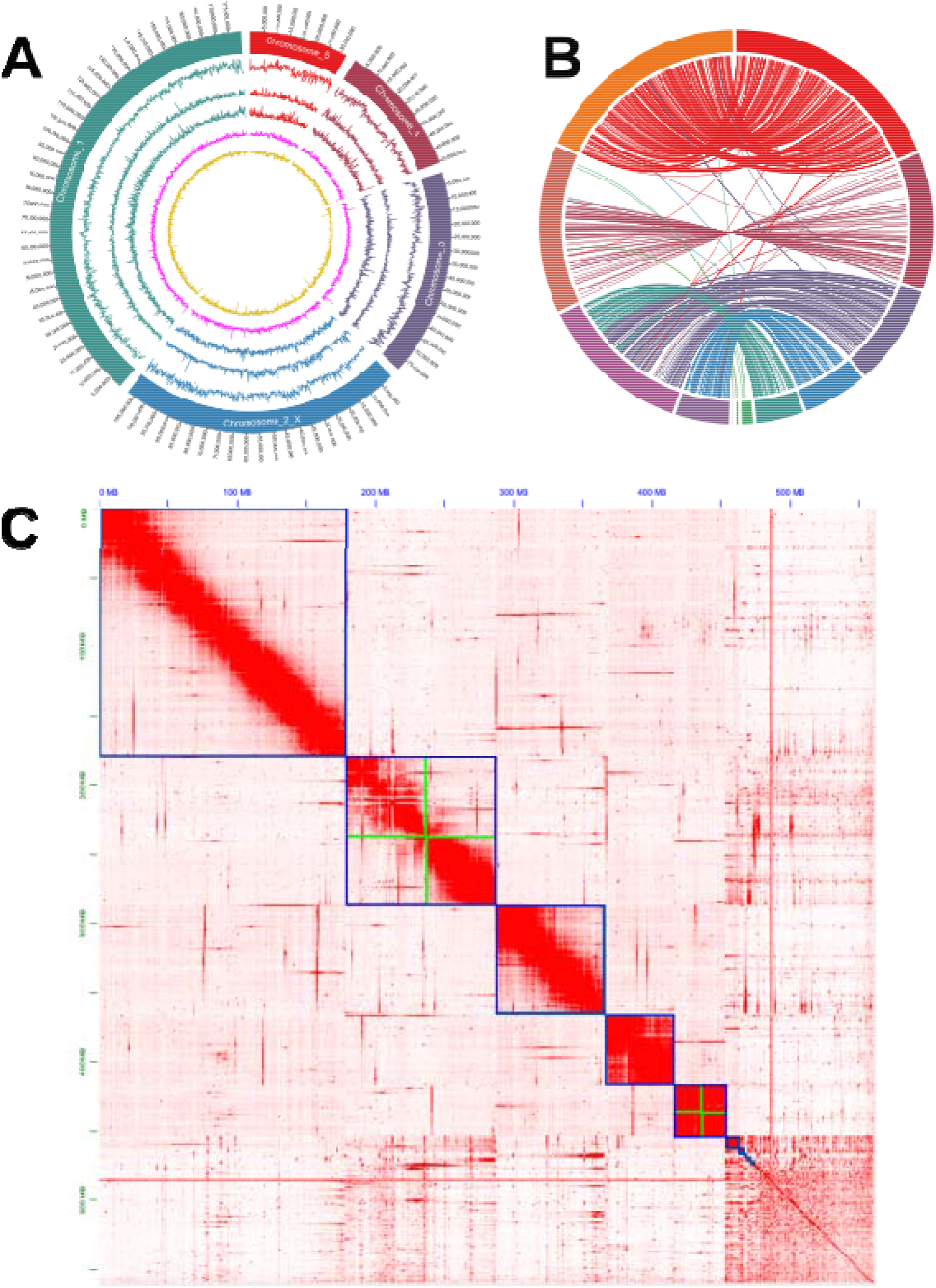
Genome structure of *Macrosiphum euphorbiae* and comparison with *Acyrthosiphon pisum*. A) Genome characteristics for *M. euphorbiae*. From outermost circle going inwards: *M. euphorbiae* chromosomes, SNP density across the genome between sequenced clonal lines, gene density, repeat density, GC%, AT%. All tracks calculated in 100kb sliding windows. B) Shared single-copy BUSCO gene synteny between *M. euphorbiae* (right hemisphere) and *A. pisum* (left hemisphere). C) Hi-C heatmap demonstrating the 5 chromosomal blocks of *M. euphorbiae*. Blue lines denote chromosomes, while green lines denote where contigs were manually appended based on Hi-C linkage information. The plot also highlights the large amount of remaining debris sequence.

The BRAKER annotation was required for phylogenetic analysis. BRAKER2 identified 31,074 protein coding genes (insecta_odb10 BUSCO: Complete: 94.2% [Single-copy: 88.9%, Duplicated: 5.3%], Fragmented: 1.5%, Missing: 4.3%, n:1367), of which 16,813 were functionally annotated using Interproscan. The phylogenetic tree based on 328 conserved single copy orthologues identifies *A. pisum* as the closest sequenced relative at chromosomal level (figure 2). Analysis of genome synteny between *M. euphorbiae* and *A. pisum* (figure 1B) shows large areas of conserved chromosome structure and macro-synteny. Lack of synteny at the end of the *A. pisum* chromosome 2 is likely attributed to rDNA arrays located in sub-telomeric regions (Blackman & Spence, 1996), with a similar gap in synteny observed between *R. maidis* and *A. pisum* (Li *et al*., 2020).

**Figure 2.**
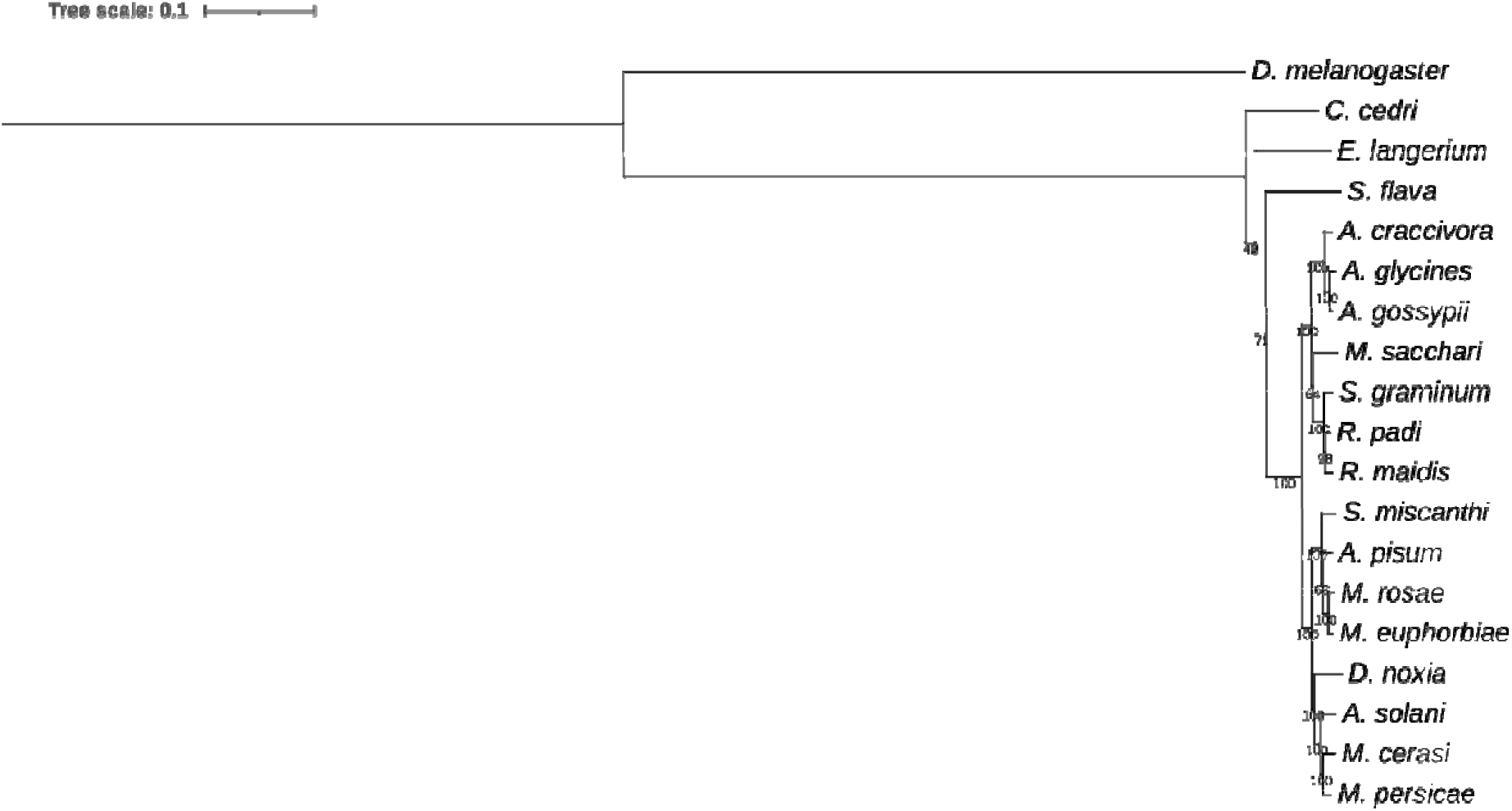
Phylogenetic analysis of aphid species based on 328 shared single copy orthologues across all species using *D. melanogaster* as an outgroup. The tree was created using a maximum-likelihood method with 1000 bootstraps. Node numbers indicate support values. Scale bar indicates amino acid substitutions per site. JTT+I+G4 was the amino acid substitution

### Comparison of Population structure at microsatellite and whole genome level

Genetic relatedness was compared between aphid clonal lines collected from geographically distinct areas over a three-year period using microsatellite and whole genome sequencing. In total 132 individual aphids were genotyped at four microsatellite positions to classify aphids into genotypes described previously (Clarke *et al*., 2017). Genotype frequency is summarised in supplementary table 2. Chi-squared analysis showed genotype proportions were not equally represented in the tested populations (*χ* ^2^ = 377.45, df = 17, P < 0.001). The majority of clones were green, with genotype 2 being the most common (n = 45/132), followed by genotype 3 (n = 34/132). Genotype 6 (n = 11/132) and genotype 7 (n = 5/132) were less abundant, as was the parasitoid-resistant genotype 1 (n = 6/132).

**Table 2.** Characteristics of aphid sampling sites for the potato aphid *Macrosiphum euphorbiae* in the three years of study.

| *Area* | *Type* | *Map key* | *Years visited* |
| --- | --- | --- | --- |
| Merseyside | plot | 1 | 2016, 2017 |
|  | plot | 2 | 2017 |
|  | plot | 3 | 2016, 2017, 2018 |
|  | plot | 4 | 2018 |
|  | farm | 5 | 2016, 2017 |
| Tayside | plot | 6 | 2016, 2017, 2018 |
|  | farm | 7 | 2016, 2017, 2018 |
| Perth | plot | 8 | 2016, 2018 |
| Fife | farm | 9 | 2016 |
|  | farm | 10 | 2016 |
|  | farm | 11 | 2016 |
| Angus | farm | 12 | 2016 |
|  | farm | 13 | 2016 |
|  | farm | 14 | 2016, 2018 |

Principal component analysis (PCA) based on these microsatellite loci demonstrated that genotypes 2 and 3 are the most distantly related (figure 3A). For green clonal lines, 12 lineages could not be assigned to a previously characterized genotype. Genotypes 4 and 5 detected in previous study (Clarke et al., 2017) were not observed over three years of sampling. Pink clones accounted for 20 sampled individuals. Within pink clones, three clusters were observed (assigned to genotypes p1, p2 and p7), while the remaining four clones could not be grouped. GLM analysis of count data including geographic location, year and site type (plot or commercial farm) indicated significantly higher abundance of genotypes 2 and 3 compared with the reference genotype 1 (see supplementary table 3). Site type was a significant factor in genotype distribution driven by the lack of genotype 3 (n = 0) found at commercial farms as this genotype was only found at plot (garden/allotment) settings (n = 34).

**Figure 3.**
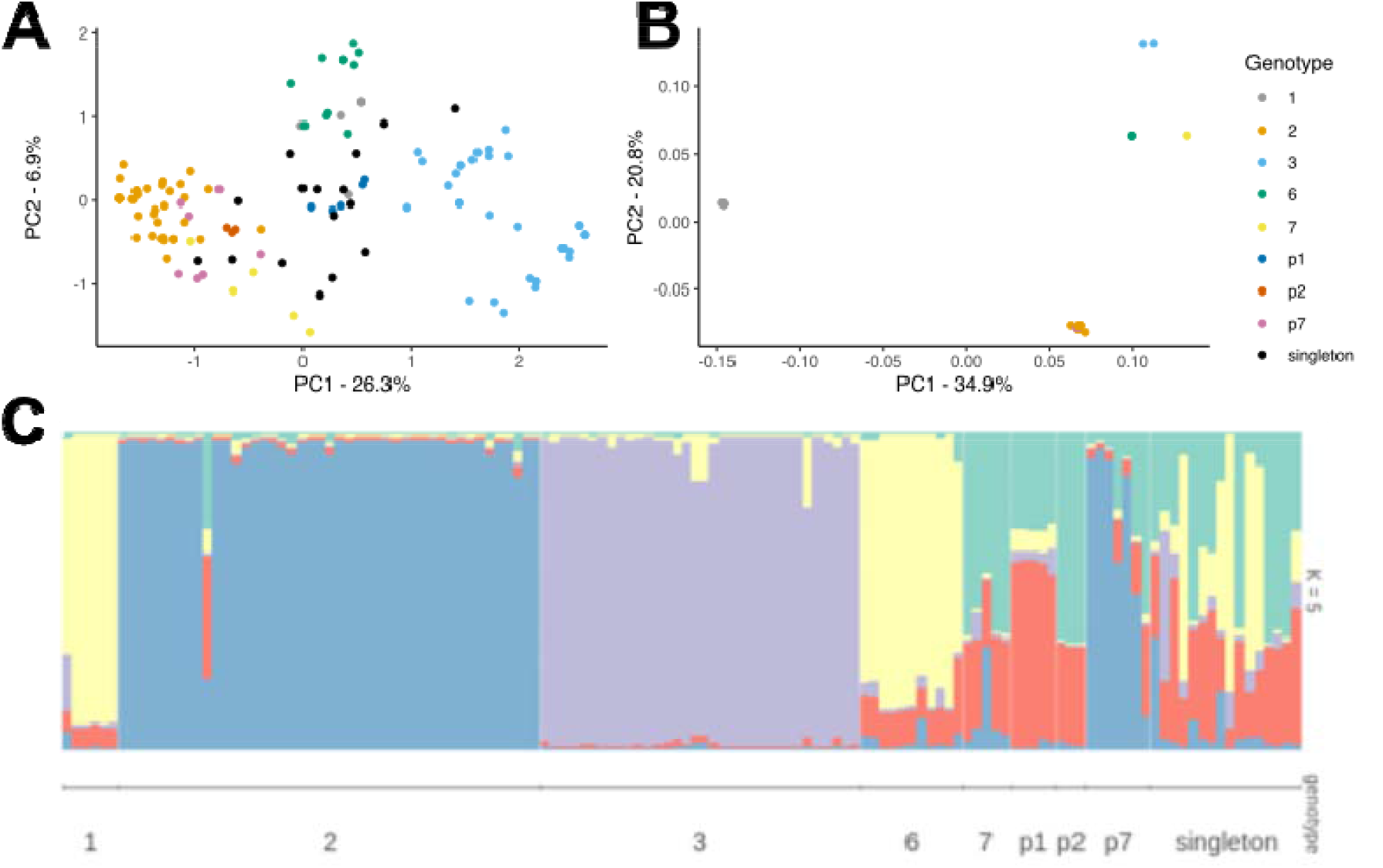
A) PCA analysis of *M. euphorbiae* individuals based on microsatellite loci. B) PCA analysis of *M. euphorbiae* individuals based on whole genome sequencing and SNP frequencies. C) *M. euphorbiae* population structure generated via STRUCTURE analysis using microsatellite data.

We observed the same genotypes over the three years of this study. Within sequenced genotypes represented by multiple clonal lines, we observe little divergence in PCA analysis. The occurrence of the same genotypes year-after-year could also indicate a lack of sexual activity in UK populations of the potato aphid, and the maintenance of asexuality over winter. As established during genome assembly, divergence between the chromosome pairs is potentially high, due to the initial lack of genome collapsing by Canu and the final 86 Mbp of sequence not being scaffolded with Hi-C. One explanation could be that these clones have hybrid origins (i.e. between ancestral aphid species), which has also been suggested to have occurred in another aphid species, *Rhopalosiphum padi* (Delmotte *et al*., 2001; Delmotte *et al*., 2002), although further analyses would be required to assess this theory. This is also summarized in Jaron *et al*. (2021), where they suggest parthenogenetic organisms with high genomic heterozygosity are a result of hybrid origin. Hybridization has been posited for genotypes of *M. persicae* that are obligate parthenogens, show high allelic divergence, and are commonly detected in populations across Australia (Vorburger et al., 2003). This dominance is similar to what we see here with *M. euphorbiae* genotypes 2 and 3.

STRUCTURE was used to assess the presence of shared genetic material between genotypes of *M. euphorbiae* using the microsatellite data. STRUCTURE and STRUCTURE HARVESTER predicted the number of population clusters (K) as 5, as denoted by the highest delta K value (supplementary figure 1). STRUCTURE output supported the PCA analysis, showing that green genotypes 1, 2, 3 and 6 formed their own clusters, as well as pink genotypes p1, p2 and p7, as can be demonstrated through uniformity in colour within each genotype and distinct colours between genotypes (figure 3C). Genetic clustering is not observed at the geographic level, suggesting a lack of recombination and predominance of asexual reproduction in UK populations of *M. euphorbiae*. As K is not equal to the number of observed distinct clonal genotypes, it could indicate genetic overlap in genotypes, or be a consequence of microsatellite data not providing enough resolution to classify genotypes correctly. The genotype counts here agree with previous studies where small numbers of *M. euphorbiae* clonal genotypes collected from potato have been identified year-after-year, suggesting persistence as obligate asexual morphs. In a study of potato aphid genotype fitness, Clarke *et al*., (2017) used 19 clonal lineages of Potato aphid, each belonging to one of seven distinct genotypes, five of which were detected in the present study (genotypes 1, 2, 3, 6, 7).

To measure genetic relatedness between clonal lineages at the genome level, a total of 3,129,782 SNP variants were retained from GATK SNP calling across all clones, of which 73,613 were retained for PCA. As previously observed through microsatellite analysis of field-collected aphid samples, most potato aphids were assigned into a known genotype group. Using a small number of microsatellite positions provides enough evidence to deduce these genotypes, although strict clonality is visualised at a higher resolution through variant analysis and clustering of individuals based on SNPs (figure 3B). Aphids of genotype 1, 3, 6 and p7 cluster away from other genotypes. Aphids belonging to the most common genotype 2 exhibit some SNP divergence based on slight separation of data points within the PCA cluster. Genotypes 2 and p7 appear the most closely related based on PCA clustering.

From 2017 and 2018, 16 samples were not placed within any previously described genotype based on microsatellite values. The increasing speed of, and volume of data generated by, next generation sequencing and the reducing cost of whole genome sequencing means that it becomes more beneficial to genotype individuals using genome wide SNP variant analysis, rather than focussing on a small set of variable length microsatellite loci via PCR (Zimmerman et al., 2020).The extra information generated at genome level can also be useful for interrogating genomes/pan-genomes for observable traits within populations (Singh *et al*., 2021).

### Possible gene functions attributing to aphid genotype and phenotype

GO term enrichment was used to identify any overrepresentation of gene functions with SNP variants. We were interested in identifying genes with sensory functions, which might indicate differences in host plant range that could underlie geographic differences in genotype abundance, and detoxification pathways, which might underpin resistance to parasitism in the resistant genotype. Amongst GO terms relating to “Biological Processes”, “Molecular Function”and “Cellular Component”, there were 65, 99 and 2 terms showing enrichment, respectively (full GO enrichment results available in supplementary table 4). Three GO terms of interest belonging to pathways involved in odorant and olfactory sensation, were identified (table 1). In addition, genes involved in detoxification pathways were identified (table 1), and have previously been linked to changes or broadening of insect host range through the ability to detoxify a broader spectrum of plant metabolites (Ramsey *et al*., 2010).

Common genotypes of *M. euphorbiae* demonstrate genotype-specific traits. For example, genotype 1 demonstrates innate resistance to parasitism by the hymenopterous parasitoid wasp *Aphidius ervi* (Clarke et al., 2017), and tends to exhibit moderate levels of esterase activity, which can contribute towards insecticide resistance (Clarke et al., 2018). Genotypes 1, 2 and 3 have been shown to differ in their ability to colonise wild and domesticated *Solanum* species, with genotype 2 faring better than genotypes 1 and 3 on the wild species *S. berthaultii* (Karley *et al*., 2018). This might imply differences between aphid genotypes in their host plant range, which has been previously observed in other aphid species (Via, 1991; Ferrari et al., 2006). Genome wide association studies (GWAS) are often useful for identifying genetic markers for these traits within populations (Weetman *et al*., 2018; Wan et al., 2019). This method is troublesome to use for *M. euphorbiae* due to its clonal nature, lack of intra-genotype genetic variability and the small number of clonal lineages observed, where sequencing hundreds of individuals may be akin to sequencing the same lineages multiple times. Therefore, the power to identify phenotype-linked variants is low, as GWAS will also identify many non-relevant variants as being linked to a measured phenotype. While we can use called variants here to identify potentially interesting genes linked to the fitness of *M. euphorbiae* genotypes, it is very difficult to link specific traits to underlying genetic markers or functional genes.

## Conclusions

Characteristics of the *M. euphorbiae* genome and its mode of reproduction provide an insight into the dynamics of an apparent asexually reproducing pest, with persistence of asexual reproduction being maintained. It is unclear whether UK Potato aphids have lost the ability to undergo sexual reproduction, or whether the UK climate in the autumn does not promote the morphological change to sexual morphs (Sandrock *et al*., 2011). Further work may involve whole-genome comparison against North American sexual-form *M. euphorbiae* lineages to help elucidate the mechanisms behind the switch to strict asexuality. For UK populations on potato crops, reasons for the dominance of specific genotypes, and whether this fluctuates, remains undetermined. Here we suggest gene functions that could be responsible for this distribution, specifically regarding host range. Further work is required to uncover the causes behind genotype 2 dominance, being unremarkable compared to other genotypes. Genotype 1 existing at a low frequency, having beneficial traits of parasitism resistance and potentially a degree of insecticide tolerance, suggests it may incur other fitness consequences that reduces its persistence.

The information in this study is useful in tailoring control measures specific for the potato aphid, exploiting differences between prominent genotypes. A scenario where we benefit from this analysis could be the controlled use of insecticides, specifically where genetic differences linked to variation in insecticide tolerance between genotypes are identified, which may inhibit efficacy of treatment. Another use for this genomic resource could also be the development of novel targets for RNAi (Christiaens *et al*., 2020).

*M. euphorbiae* remains an important pest in vectoring multiple plant pathogens over a variety of crops. While still presenting some challenges regarding genome assembly and annotation, the work here not only provides a near-chromosome level assembly for *M. euphorbiae*, but also provides insights into the genetic structure and diversity of *M. euphorbiae* in UK populations. Due to the clonal nature of *M. euphorbiae* in the UK, the sequence data generated in this study likely represents much of the genetic sequence of UK *M. euphorbiae* on potato crops.

## Supporting information

Supplementary tables and data sources

## Data availability and supplementary files

Read files and assemblies can be found under the ENA project accession PRJEB55422. A full description of read file accessions and data sources for analyses can found in supplementary tables 5 and 6.

## Acknowledgments

We would like to the thank Margaret Hughes (Centre for Genomic Research (CGR), University of Liverpool (UK)) for PacBio library prep and sequencing, as well as Charlotte Nelson (CGR) and Anita Lucaci (CGR) for Illumina library prep and sequencing respectively. We would also like to thank Marta Maluk, Desiré Macheda and Carolyn Mitchell at the James Hutton Institute (Dundee, UK) for help in aphid sampling and PCR genotyping. The project was funded through the Biotechnology and Biological Sciences Research Council (BBSRC) iCASE award and James Hutton Limited. AJK is supported by the Strategic Research Programme funded by the Scottish Government’s Rural and Environment Science and Analytical Services (RESAS).

## Methods

### Aphid sampling and stats

Over three field seasons, potato aphids were collected from private allotments and commercial farms around the Merseyside (supplementary figure 2) area as well as the Tayside, Perthshire and Fife areas of Scotland (supplementary figure 3) (table 2). For commercial farms, between three and five individuals were collected at loci separated by at least 50 paces to avoid sampling aphids that had reproduced clonally in the locality. This approach was not possible at allotment sites, and instead individuals were collected from different allotment plots. In 2016 and 2017, collected aphids were kept at constant conditions of 16h:8h L:D, 20 °C ± 1 and 70% RH until determined to be healthy and free of parasitoids and fungal pathogens. Aphids were maintained under these conditions during clonal culture and individuals were sampled from live cultures. In 2018, aphids were immediately frozen after collection and stored at -80 °C until analysis.

### Aphid DNA extraction and genotyping

Between 1-5 aphids of each clonal lineage were flash frozen with liquid nitrogen and homogenised in a 1.5 mL tube with a pestle. For samples collected in 2016 and 2018, DNA was extracted using the Qiagen Dneasy Blood and Tissue Kit (Hilden, Germany). In 2017, the Zymo gDNA miniprep Kit (California, USA) was used. In both cases, DNA was extracted following the manufacturer’s protocol. DNA quality and quantity was assessed using Nanodrop (ThermoFisher, Massachusetts, USA).

Genotyping was performed with seven microsatellite loci for samples collected in 2016 (*Me1, Me5, Me7, Me9, Me10, Me11* and *Me13*). Primer design is outlined in Raboudi et al. (2005) and supplementary table 7 (PCR protocol outlined in supplementary tables 8 and 9). Only four marker sites (*Me1, Me5, Me9 and Me10*) were used for samples collected in 2017 and 2018 to streamline analysis and reduce costs, as data from 2016 samples showed these marker sites were sufficient to differentiate the clonal lineages. Microsatellite loci were amplified via PCR. Lengths of the resulting PCR amplicons were measured via capillary electrophoresis on a 3730 DNA Analyzer (ThermoFisher, Massachusetts, USA) using the GeneRox 500 internal standard. Microsatellite lengths were assessed using the Genemapper 5 software.

### Statistical analyses on genotype frequencies

Analyses were performed using R (v3.2.2). Graphical output was generated using R and ggplot2 (v3.1.0) (Wickham, 2016). Chi-squared analysis was used to assess genotype distribution over three years of sampling. GLM analysis of count data with Poisson distribution was used to assess the effect of year, sampling location and site type on observed genotype frequencies. Adegenet (v2.1.1) (Jombart, 2008) in R was used to generate PCA plots with frequencies of amplicon lengths at each microsatellite locus (*Me1, Me5, Me9* and *Me10*) using default settings and visualized in ggplot2.

### Structure analysis

Population structure based on four microsatellite loci available for all samples (*Me1, Me5, Me9* and *Me10*) was generated using the Bayesian clustering algorithm STRUCTURE (v2.3.4) (Pritchard *et al*., 2000). STRUCTURE analysis demonstrates genetic relationships between genotypes and evidence of any recombination before the switch to parthenogenesis. Parameters include a Burnin period of 10,000 with 50,000 MCMC replicates after Burnin. For K of 3 to 24 (with 24 being the theoretical maximum based on unique genotypes described here), 10 replicates were performed, where K is the number of distinct genetic groups. STRUCTURE output was uploaded to STRUCTURE HARVESTER (web v0.6.94) (Earl & von Holdt, 2012), an online tool to predict the best fitting value of K produced from STRUCTURE using the Evanno delta K method (Evanno et al., 2005). STRUCTURE output for K= 5 was visualised using the R package Pophelper (v2.3.1) (Francis, 2017).

### High Molecular Weight (HMW) DNA extraction

An expanded version of this protocol is outlined in supplementary methods. Briefly, HMW DNA was extracted from 50 adults of genotype 1 (MW16/67; supplementary table 13) *M. euphorbiae* using a phenol-chloroform method and phase-lock separation gel (Quantabio, Massachutsetts, USA), where phase-separating gel allows no pipetting and results in much less shearing of DNA molecules. DNA quality was assessed using 1 μl of DNA (diluted 1 in 10 in nuclease-free water), with purity measured using Nanodrop and quantity estimated using Qubit.

### Short read DNA sequencing

For initial sequencing of six genotype 1 clonal lines, library preparation was carried out by the Centre for Genomic Research (CGR) in the University of Liverpool. TruSeq PCR free libraries (2x150 bp) with a 550 bp insert were generated for the six genotype 1 aphid samples. Libraries were sequenced on a single lane of the Illumina HiSeq 4000. A further 10 TruSeq PCR free libraries were generated consisting of other potato aphid genotypes for sequencing on the Illumina 4000, as well as two TruSeq PCR free libraries for samples MW16/48 and AK13/30 (supplementary table 13) (2x300 bp) for the Illumina HiSeq 2500 (rapid run mode). All reads were trimmed for Illumina adapters using Cutadapt (v1.2.1) (Martin, 2011) followed by further trimming and quality filtering with Sickle (“-q 20”and “-l 20”enabled) (v1.200) (Joshi & Fass, 2011).

### Long read DNA sequencing

High molecular weight DNA library preparation and sequencing was performed by CGR, with resulting libraries sequenced on the PacBio Sequel using v1.2.1 chemistry. DNA was sheared to 20 Kb. DNA was sequenced over 10 SMRT cells, providing 60 Gb of data in total. The same high molecular weight DNA was used to prepare a single 10x chromium library and sequenced on a lane of the Illumina HiSeq 2500 (paired-end sequencing; 2x150 bp). Library preparation and sequencing for Hi-C libraries was performed by Dovetail genomics (California, USA).

### Contig assembly

Initial draft assemblies were generated using Canu (v1.4) (Koren *et al*., 2017) using default parameters and were provided with a genome estimate size of 530 Mb (Wenger et al., 2017). Three iterations of arrow were performed to polish the Canu assembly prior to any downstream analysis. For each iteration, the Canu reference was uploaded to the smrtlink (v4.0.0) portal and provided with all PacBio reads used in the initial assembly. Alternative haplotypes were removed using HaploMerger2 (v20161205) (Huang et al., 2017), a tool designed to provide a haploid assembly from a diploid sequence through self-alignment with LASTZ (Harris, 2007). Haplomerger2 also requires the use of a custom script (lastz_D_wrapper.pl) to generate a score matrix at 95% identity (Huang et al., 2017). Haplomerger2 was used until genome size was reached of approximately 530 Mb based on Wenger et al. (2017) and GenomeScope analysis.

### Scaffolding and chromosome arrangement

Contig orientation and scaffolding was performed first with 10x linked reads using arcs (v1.0.1) (Yeo *et al*., 2018). 10x GemCode barcodes were manually appended to fastq headers and mapped using BWA-MEM (v0.7.5a) (Li & Durbin, 2009) and the resulting bam file provided to arcs. Further scaffolding with Hi-C data was performed by Dovetail genomics using their HiRise assembly platform (California, USA). Hi-C contact maps were generated using Juicer (v1.5) and visualised using Juicebox (v1.8.9) (Durand et al., 2016). The final assembly was polished with ntedit (v1.3.0) (Warren *et al*., 2019). For the predicted X chromosome and chromosome 5, manual assembly was required using JBAT.

### Buchnera assembly

A hybrid-assembly approach was taken for the Buchnera genome. For Illumina reads for the clonal line MW16/67, reads were taxonomically assigned using kraken2 (v2.1.2) (Wood *et al*., 2019). Buchnera assigned reads were retrieved using extract_kraken_reads.py with ‘— taxid 9’, the taxon ID for *Buchnera*. Draft assembly of filtered reads was generated using spades (v3.15.3) (Prjibelski *et al*., 2020) with the options ‘—cov-cutoff 100 —isolate’. *Buchnera* contigs assembled from the PacBio assembly were then identified from Blobtools analysis, with these contigs then used to scaffold the spades assembly using RagTag (v2.1.0) (Alonge et al., 2022). The *Buchnera* contig was identified based on its length (645.9 Kbp). Plasmid sequences for pTrp and pLeu were identified through Blast (v2.12.0+) (Camacho et al., 2009). The *Buchnera* genome and its plasmids can be found under the ENA accession ERS14404677.

### RNA extraction and ONT sequencing

RNA was extracted from 200 individual aphids from clonal culture MW16/67 (genotype 1), consisting of a mixture of instars. Aphids were divided over four 2 mL Eppendorf tubes (50 aphids each). Aphids were flash frozen in liquid nitrogen and homogenised using a pestle. Cells were re-suspended in 500 μl TRI-reagent (Sigma-Aldrich, Missouri, USA) and left at room temperature for 5 minutes. RNA from the cell solution was isolated using the Directzol RNA MiniPrep kit (Zymo-research, California, USA) following the procedure outlined in the kit, as well as performing the optional DNase step. mRNA isolation was performed using Dyna-beads (Thermofisher, Massachusetts, USA) using the procedure outlined within the kit. Qubit and Nanodrop values were obtained throughout the process. As mRNA quantities were low after isolation with Dyna-beads, it was decided that integrity would be measured after cDNA synthesis to ensure as much material as possible was kept for MinION sequencing.

For transcriptome sequencing, 250 ng of purified mRNA was used as input for library preparation using Direct cDNA sequencing kit (SQK-DCS109) (Oxford Nanopore Technologies (ONT) Oxford, UK). The kit Direct cDNA requires no PCR to avoid introducing PCR bias. Library preparation was performed as outlined in the accompanying protocol, with a minor change. After reverse strand synthesis and adapter ligation, five separate libraries were pooled to generate as much sequencing material as possible. Pooled libraries were sequenced using two FLO-MIN106 version flow cells and MinKnow (v18.03.1). Base calling was performed using Albacore (v2.3.1) (Wick *et al*., 2019).

### Genome annotation

Draft annotation was generated using publicly available RNA-seq data from *M. euphorbiae* (Teixeira et al., 2018; NCBI Sequence Read Archive accession SRX339176) and ONT generated cDNA. Briefly, Illumina RNA seq reads were trimmed with Trimmomatic (v0.39) (Bolger *et al*., 2014), followed by mapping to the genome with STAR (v2.5.2a_modified) (Dobin *et al*., 2013) to genome softmasked using RepeatModeler (v.1.0.11) (Smit *et al*., 2008) and RepeatMasker (v4.0.7) (Smit et al., 2013). cDNA reads from ONT sequencing were mapped using minimap2 (v2.2-r424-dirty) (Li, 2018). Bam files for each read set were provided to BRAKER2 (v2.1.5) (Brůna *et al*., 2021) for automatic genome annotation. A second BRAKER2 (--ep mode enabled) was used for gene prediction based on A. pisum proteins, followed by gene model selection using TSEBRA (Gabriel et al., 2021) with default settings. The longest isoform for each gene was retained for further analyses. Predicted proteins were functionally annotated with Interproscan (v5.32-71.0) (Jones *et al*., 2014), with “—goterms”enabled.

### Phylogenetic tree generation

Gene predictions from BRAKER2 were used to place *M. euphorbiae* within the aphid phylogeny. Orthogroups consisting of single-copy orthologues across all species were identified using orthofinder (v2.5.4) (Emms & Kelly, 2019), followed by alignment and alignment trimming with mafft (v.7.487) (Nakamura *et al*., 2018) and gblocks (v0.91b) (Castresana, 2002) respectively. Alignments were concatenated and provided to modeltest-ng (v0.1.7) (Darriba *et al*., 2020) to calculate the best-fit amino acid substitution model, with the resulting model supplied to RaxML-ng (v0.6.0) (Kozlov *et al*., 2019) along with the alignment for phylogenetic tree inference. RaxML-ng was executed using 10 randomized parsimony trees, followed by 1,000 bootstrap replicates.

### Variant calling

Single nucleotide polymorphisms (SNP) were generated for analysing genetic relatedness between Potato aphid genotypes as well as gene nucleotide diversity. SNP variants were called using GATK (v3.7) (Mckenna et al., 2010). Illumina reads from each clonal line sequenced were mapped using BWA-MEM (v0.7.5a-r405) (Li & Durban, 2009). Mapped reads were re-aligned around indels using the two GATK tools ‘RealignerTargetCreator’ and ‘IndelRealigner’. ‘HaplotypeCaller’ was used to generate variant calls for each clonal line mapped to the genotype 1 reference (--GVCF enabled), followed by joint genotyping and combing of individual sample variant calls using ‘GenotypeGVCFs’. SNP variants were extracted, then filtered using ‘VariantFiltration’, using the following filter expressions; “QUAL < 0 || MQ < 40.00 || SOR > 4.000 || QD < 2.00 || FS > 60.000 || MQRankSum < - 20.000 || ReadPosRankSum < -10.000 || ReadPosRankSum > 10.000”for SNPs. Finally, variants were retained where a genotype was called for each individual.

Divergence of Potato aphid genotypes was assessed via PCA. A subset of SNPs was generated with a minimum of 5000 bp between each-other using vcftools (“—thin 5000”) (v0.1.13) (Danecek et al., 2011). Genotype clustering and MDS tables were generated through PLINK (v1.90p) (Purcell et al., 2007) (“—noweb”and “—allow-no-sex”enabled). The multidimensional scaling (MDS) table contains 10 principal components based on genetic distance between individuals using SNPs; the first two principal components, PC1 and PC2, were used for PCA. PCA variation for each component was calculated as a percentage of the sum of all eigenvalues in the generated “plink.eigenval”output for each individual component. PCA plots were generated in R (v3.4.4) (R core team, 2015).

### Identifying gene functions of interest

SNP variants (genotyped across all sequenced clonal lineages) were annotated using SnpEff (v4.3t) (Cingolani et al., 2012) for missense mutations. Genes containing missense mutations were used in GO term enrichment analyses to identify gene functions with increased genetic variation. GO enrichment was carried using TopGO (v2.38.1) (Alexa & Rahnenfuhrer) with summary statistics generated using the web tool Revigo (Supek *et al*., 2011).

