## Supplementary figures for "Genomic insights into clonal diversity in UK populations of the Potato aphid, *Macrosiphum euphorbiae*"

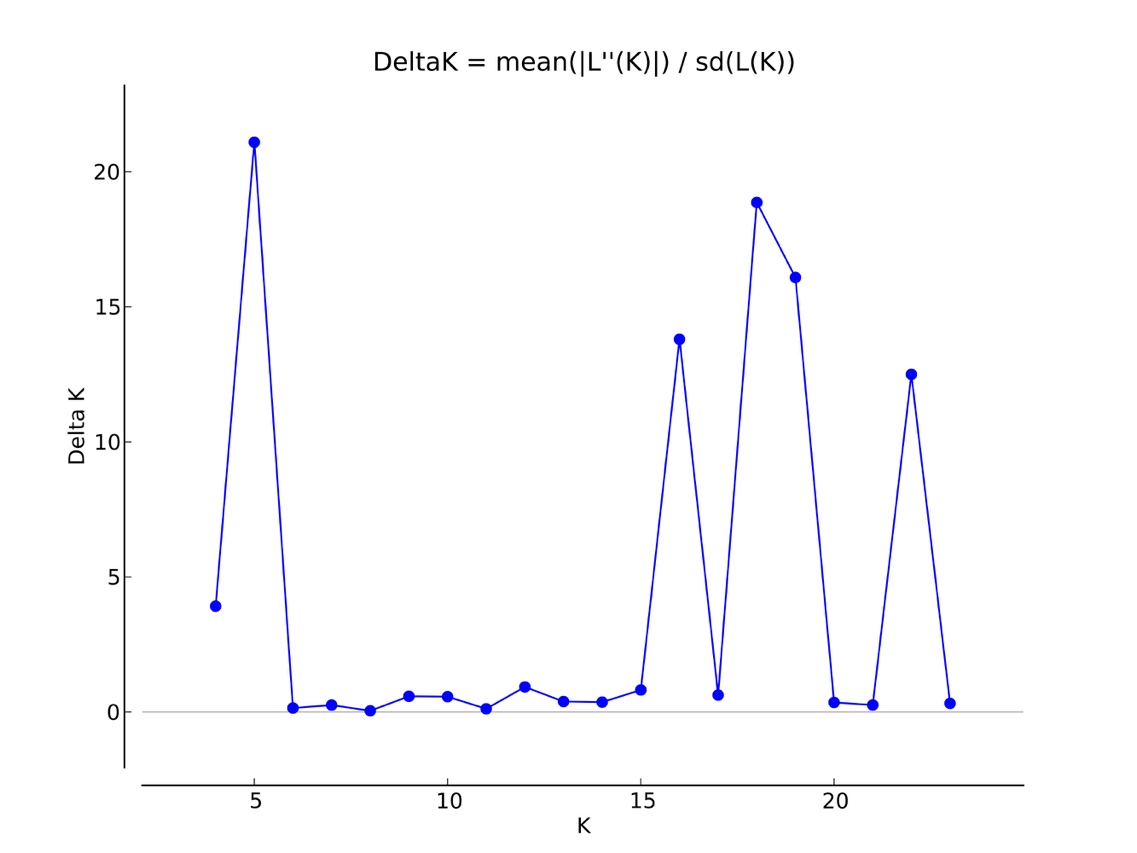

**Supplementary figure 1. Predicted value of genetic clusters (*K*) for *M. euphorbiae* based on the delta *K* method implemented in STRUCTURE HARVESTER (Earl & von Holdt, 2012).** Briefly, delta K is calculated through measuring the highest likelihood of each value of *K*, followed by assessing the variance of likelihood values between replicates within each *K.*

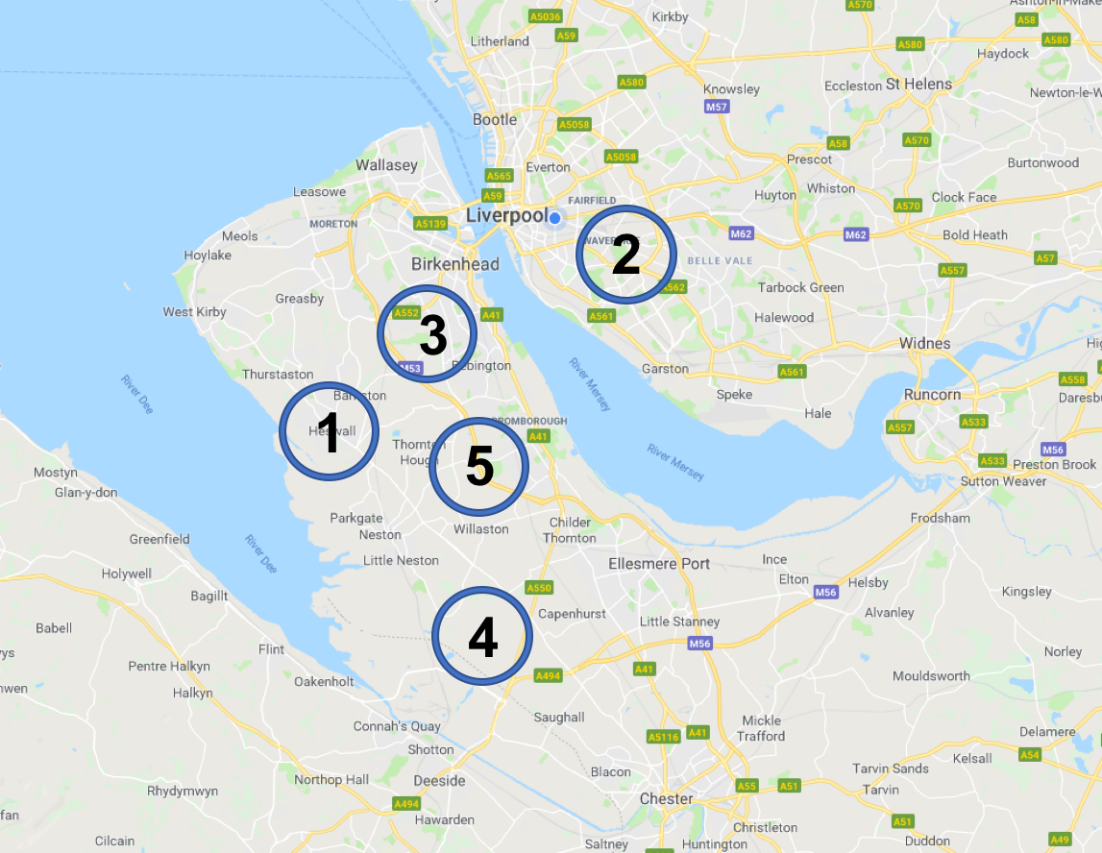

**Supplementary Figure 2. Locations of Merseyside sites used for aphid sampling.** Also see supplementary table 2.

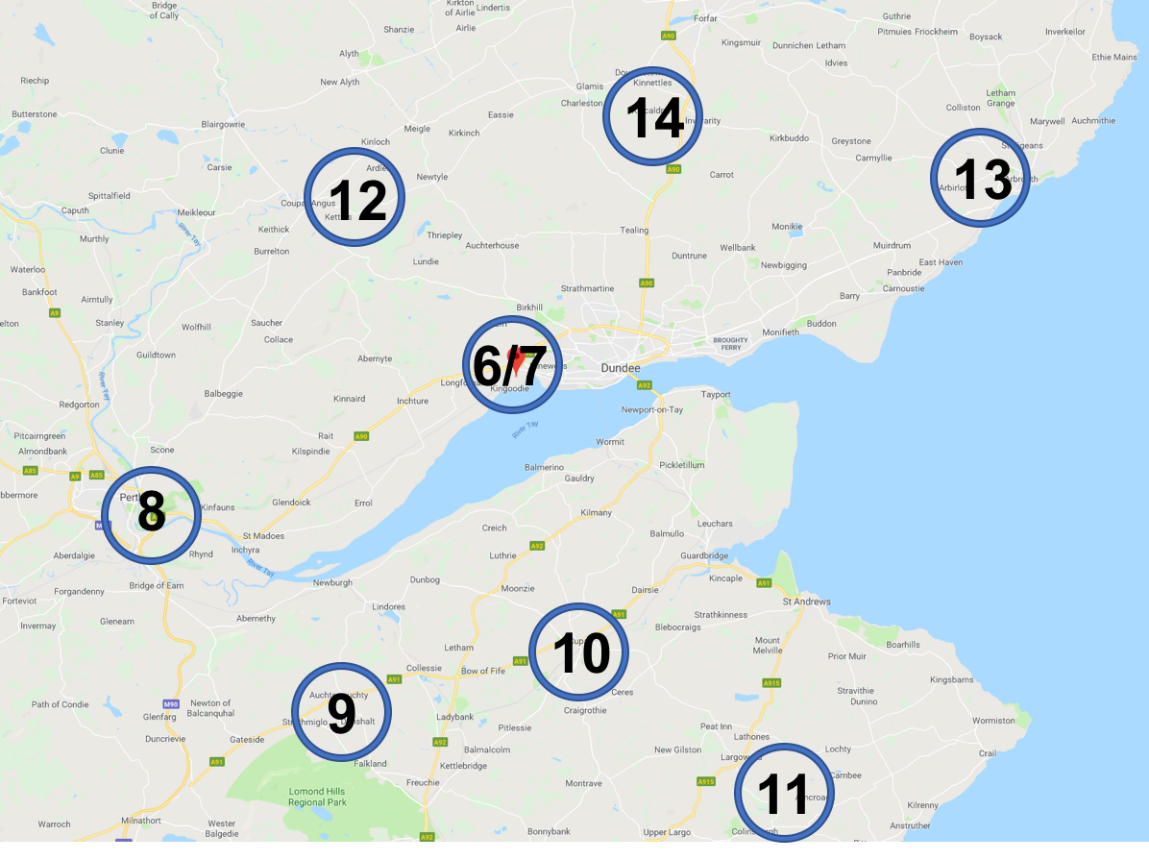

**Supplementary Figure 3. Locations of sites in Tayside, Perth and Fife used for aphid sampling.** Also see supplementary table 2.
