## Supplementary methods for "Genomic insights into clonal diversity in UK populations of the Potato aphid, *Macrosiphum euphorbiae*"

*High molecular weight DNA preparation and Long-read library sequencing*

The method described is adapted from the Quick protocol (Quick, 2018). Fifty adult aphids of each clonal line were divided between three 2 mL Eppendorf tubes. The samples were flash frozen in liquid nitrogen and homogenised using a pestle. One mL of tissue lysis buffer (10 mM Tris-Cl pH 8.0; 25 mM EDTA pH 8.0; 0.5% SDS) was added to the homogenised insect powder. The three aliquots were then pooled in a 50 mL Falcon tube with an extra 2 mL of lysis buffer. RNAse A (20 mg/mL) was added to a final concentration of 20 μg/mL, the tube slowly rotated end-over-end ten times and incubated at 37 °C for 1 hour. 25 μl Proteinase K (20 mg/ml) was then added and the solution incubated at 50 °C for 2 hours, with 10 more slow end-over-end rotations performed every hour.

The lysis solution was decanted into a 15 mL tube containing light-variety phase lock gel (Quantabio, Massachutsetts, USA), along with 5 mL of phenol solution. The tube was placed on a rotary shaker at 40 rpm for 10 minutes until a “milky” mixture had formed. The sample was centrifuged at 2,300 *g* for 12 minutes, after which the aqueous phase was poured into a new 15 mL tube containing phase lock gel. The gel is less dense than the organic phase of the phenol, resulting in a cleaner extraction and DNA which is less sheared compared to methods that require pipetting. 5 mL chloroform/isoamyl (25:1) was added and the sample placed back on the rotary shaker for 10 minutes at 40 rpm followed by centrifugation at 2300 *g* for 12 minutes. The aqueous phase was decanted into a 50 mL tube.

To precipitate the DNA, 15 mL of ice cold ethanol was added to the aqueous phase, along with 2 mL of sodium acetate (1M; pH 5.2) and mixed by careful inversion, rotating the tube gently end-over-end 10 times. The ‘jelly’-like DNA precipitate was then “pulled” from the solution using a glass Pasteur pipette with a hooked end, made by briefly heating the end up in a Bunsen burner. DNA was washed by submerging in 70% ethanol until turning opaque, then was carefully removed from the pipette into a new 2 mL tube. DNA was centrifuged in 1 mL of 70% ethanol for 1 minute at 10,000 *g*. Ethanol was removed with a pipette without disturbing the DNA pellet. The pellet was then left to air-dry at 40 °C for 10 minutes to remove residual ethanol. DNA was re-suspended without agitation in 50 μl nuclease-free water and left to slowly dissolve overnight at 4 °C. DNA quality was assessed using 1 μl of DNA (diluted 1 in 10 in nuclease-free water), with purity measured using Nanodrop and quantity estimated using Qubit.

High molecular weight DNA library preparation and sequencing was performed by CGR, with resulting libraries sequenced on the PacBio Sequel using v1.2.1 chemistry. DNA was sheared to 20 Kb. DNA was sequenced over 10 SMRT cells, providing 60 Gb of data in total. The same high molecular weight DNA was used to prepare a single 10x chromium library and sequenced on a lane of the Illumina HiSeq 2500 (paired-end sequencing; 2x150 bp). Library preparation and sequencing for Hi-C libraries was performed by Dovetail genomics (California, USA).

*Symbiont PCR*

As well as genotyping, we also partially characterised the symbiont infection status of collected aphid lines. DNA was extracted as described in the methods section ‘Aphid DNA extraction and genotyping’. Purified DNA was assessed for symbiont presence by polymerase chain reaction (PCR) and amplification of the 16S rDNA gene and 16-23 S region (specific region absent in *Buchnera*), where 16-23 S positive results indicate secondary symbiont presence. Primers specific for *Hamiltonella defensa, Serratia symbiotica* and *Regiella insecticola* are outlined in supplementary tables 10, 11 and 12*.* PCR amplicons were visualized using gel electrophoresis and compared against an *Escherichia coli* positive control*.*

No further analysis was performed on the symbiont data, however it is provided here for completeness (see supplementary table 13)

*References*

Quick, J., 2018. Ultra-long read sequencing protocol for Oxford Nanopore. *Protocols.io.* https://www.protocols.io/view/ultra-long-read-sequencing-protocol- for-oxford-nan-k88czzw?version_warning=no.
